# Online abstraction during statistical learning revealed by neural entrainment from intracranial recordings

**DOI:** 10.1101/2023.01.11.523605

**Authors:** Brynn E. Sherman, Ayman Aljishi, Kathryn N. Graves, Imran H. Quraishi, Adithya Sivaraju, Eyiyemisi C. Damisah, Nicholas B. Turk-Browne

## Abstract

We encounter the same people, places, and objects in predictable sequences and configurations. These regularities are learned efficiently by humans via statistical learning. Importantly, statistical learning creates knowledge not only of specific regularities, but also of more abstract, generalizable regularities. However, prior evidence of such abstract learning comes from post-learning behavioral tests, leaving open the question of whether abstraction occurs online during initial exposure. We address this question by measuring neural entrainment during statistical learning with intracranial recordings. Neurosurgical patients viewed a stream of scene photographs with regularities at one of two levels: In the Exemplar-level Structured condition, the same photographs appeared repeatedly in pairs. In the Category-level Structured condition, the photographs were trial-unique but their categories were paired across repetitions. In a baseline Random condition, the same photographs repeated but in a scrambled order. We measured entrainment at the frequency of individual photographs, which was expected in all conditions, but critically also at half of that frequency — the rate at which to-be-learned pairs appeared in the two structured conditions (but not the random condition). Neural entrainment to both exemplar and category pairs emerged within minutes throughout visual cortex and in frontal and temporal brain regions. Many electrode contacts were sensitive to only one level of structure, but a significant number encoded both exemplar and category regularities. These findings suggest that abstraction occurs spontaneously during statistical learning, providing insight into the brain’s unsupervised mechanisms for building flexible and robust knowledge that generalizes across input variation and conceptual hierarchies.

## Introduction

Everyday experience is highly structured and humans can learn this structure via a process known as statistical learning (***Sherman et al., 2020***). This knowledge in turn lets us generate predictions and behave more efficiently when we encounter familiar environments in the future. For example, after repeatedly traveling through a local airport, you know where to park, how to check-in, which security lines are efficient, where to stop for good food, and how the gates are arranged, all of which makes travel smoother than in a foreign airport. At the same time, beyond the specifics of your local airport, many features of your experience reflect general properties of air travel that generalize to most or all other airports, including ground transportation, security, boarding, baggage, etc., meaning that experienced travelers can still intuit what to do even in a new airport.

Prior behavioral studies have shown that statistical learning supports this kind of abstraction (***Brady and Oliva, 2008***; ***Otsuka et al., 2013***; ***Emberson and Rubinstein, 2016***; ***Jun and Chong, 2018***; ***Luo and Zhao, 2018***; ***Jung et al., 2021***). A common design in such studies is to expose participants to a sequence of images with regularities at the category level (e.g., images of beaches always followed by images of canyons); this differs from standard studies of statistical learning in which the regularities exist at the level of particular exemplar images that repeat in pairs or triplets. Evidence for category-level statistical learning is assessed offline in a behavioral test after sequence exposure, for example, by asking participants to rate the familiarity of a category pair to which they were exposed (e.g., beach -> canyon) vs. a foil (e.g., beach -> farm, where farm was a category in another pair). The categories in these test items are often represented by novel exemplars or category labels, such that they can only be discriminated if the participants abstracted categorical regularities that they can generalize to these novel stimuli.

These prior studies usefully demonstrated that statistical learning supports abstraction, but the use of offline tests limits insight into learning process itself. Specifically, it is unclear how and when participants form these abstract representations, and critically whether this occurs *during* learning at all. Rather, it is possible that participants learn the specific regularities to which they were exposed and only at test do they abstract these regularities to novel exemplars or labels through analogy or inference. For example, if exposed to a pair of exemplars during learning (e.g., beach1 -> canyon1), participants may exhibit familiarity or discrimination for new exemplars of the same categories at test (e.g., beach2 -> canyon2) either (1) because they had already abstracted a general category relation online during exposure that is ready to be applied, or (2) because no abstraction occurred in advance and they instead retrieve specific learned pairs and infer that the right answer will preserve the same category relation. This theoretical distinction of whether inference occurs during encoding or retrieval has been examined in other forms of learning and memory (***Preston and Eichenbaum, 2013***; ***Zhou et al., 2021***). A prior study from our lab provided some tentative behavioral evidence that abstraction might occur online during statistical learning (***Sherman and Turk-Browne, 2020***), which prompted us to conduct a targeted study to measure online abstraction during statistical learning of category-level regularities more directly.

For this purpose, we adopted a technique known as neural entrainment (or frequency tagging) that has found recent success in tracking statistical learning of auditory and visual regularities (***Ding et al., 2016***; ***Batterink and Paller, 2017***; ***Batterink, 2020***; ***Choi et al., 2020***; ***Henin et al., 2021***). This electroencephalography (EEG) based method capitalizes on the fact that brain oscillations in sensory regions can exhibit phase locking, or entrainment, at the frequency of onset of rhythmic stimuli (***Norcia et al., 2015***; ***Bauer et al., 2020***). The presence of such entrainment can be used to detect whether and where in the brain the stimuli are processed, including when multiple stimuli are presented at different frequencies (***Nozaradan et al., 2011***; ***Störmer and Alvarez, 2014***; ***Ding et al., 2016***). Indeed, statistical learning studies have found neural entrainment not only at the frequency of individual stimuli, but also to the frequency of learned groupings of multiple stimuli (***Henin et al., 2021***), despite no explicit segmentation cues indicating these groupings.

For example, in a visual stream in which certain scenes follow each other with high transition prob-ability constituting pairs, neural entrainment is expected at the frequency of individual scenes, reflecting visual-evoked responses, but also at half of that frequency, reflecting the rate of learned pairs. Critically, this provides a measure of learning because the pairs only exist in the minds of participants who extracted them based on statistical regularities across repetitions (again, there is no explicit timing, instruction, or other cue in the stimuli about the existence of pairs). Because entrainment to learned regularities is measured incidentally while participants are passively exposed to the stream, this method provides a continuous *online* measure of statistical learning not readily available in behavior. Beyond being a sensitive online measure of learning, neural entrainment can also provide mechanistic insight into the learning process. For example, it can help elucidate the timecourse of statistical learning by quantifying how much exposure is required for entrainment to emerge. Moreover, with the higher spatial resolution and coverage of deep-brain structures provided by intracranial EEG, it is also possible to localize statistical learning effects in the brain.

Prior studies of statistical learning with neural entrainment employed stimuli that were identical across repetitions, leaving open the question of whether abstraction occurs online during statistical learning. Thus, we combine, for the first time, the method of neural entrainment as an online measure with a task design optimized for evaluating categorical abstraction during statistical learning. This task builds on our recent intracranial EEG (iEEG) study that conflated regularities at the exemplar and category level (***Sherman et al., 2022***). Here we evaluate these two levels of abstraction separately in distinct conditions (relative to a random baseline condition), allowing us to quantify neural entrainment online during exemplar-level and category-level statistical learning.

Across task runs we manipulated the nature of regularities in a sequence of scene images (Figure 1): in Category-level Structured runs, each image appeared once such that regularities could exist only at the level of categories (e.g., category A -> category B); this differed from Exemplar-level Structured runs with repeating images that contained regularities at the level of individual exemplars (e.g., scene A -> scene B); both of these Structured runs with regularities were compared to a Random run in which images repeated without any regularities in their temporal order. Patients were not informed about these different conditions or about the presence of regularities, and they learned them incidentally through passive exposure. By capitalizing on the spatial and temporal resolution of intracranial EEG, we tracked statistical learning of exemplar and category regularities across the brain, providing insight into how, when, and where abstraction occurs.

**Figure 1.**
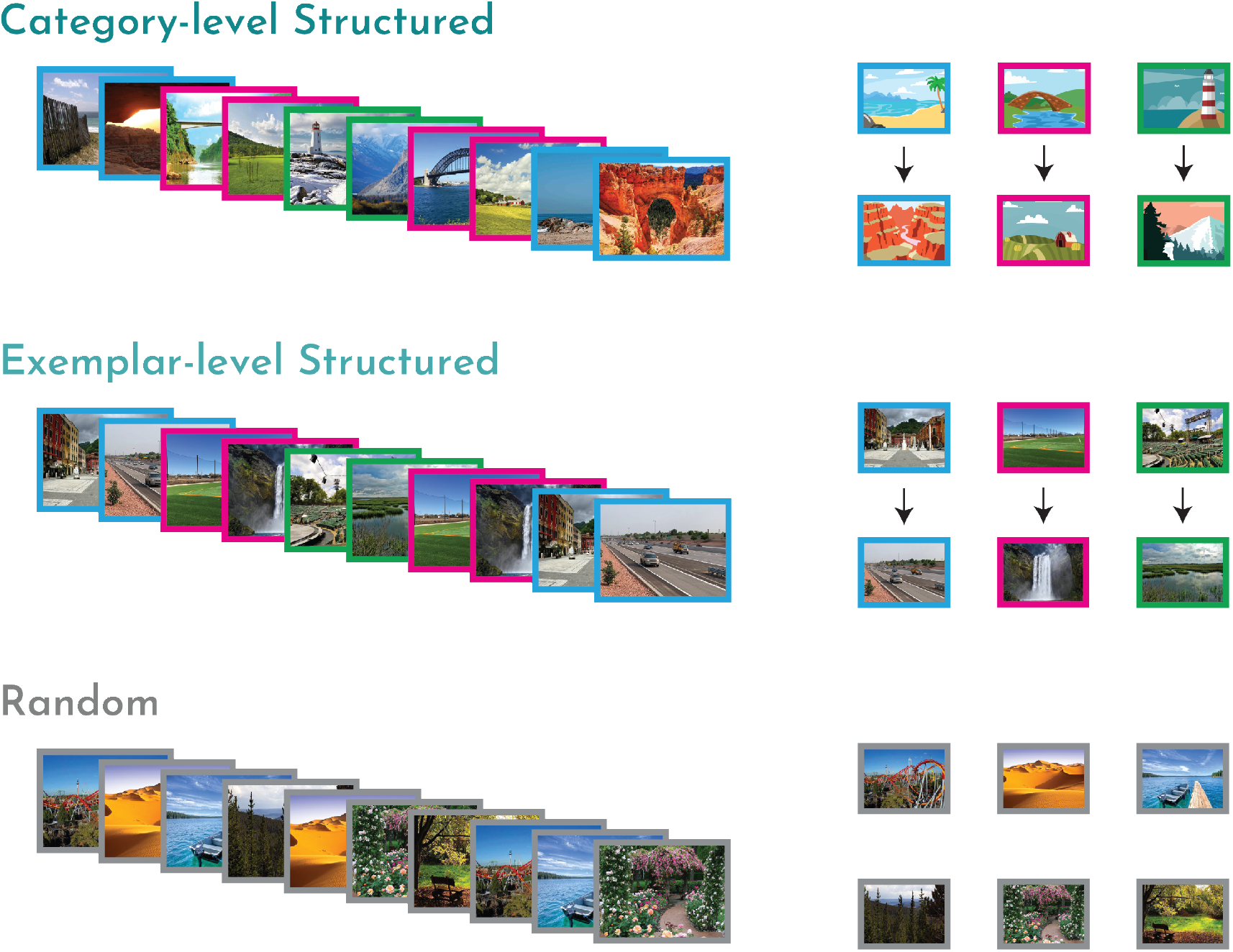
Task design and experimental conditions. Participants viewed a rapid stream of scene images (left), with varying levels of temporal structure (right). In the Category-level Structured condition (top), participants encountered a series of trial-unique scene images, drawn from six scene categories. Scene categories were temporally paired (three pairs of two categories), such that an image from one category (e.g., beach) was always followed by an image from another category (e.g., canyon). In the Exemplar-level Structured condition (middle), participants encountered a total of six scene images that appeared in temporal pairs. In the Random control condition (bottom), participants again encountered six (novel) scene images but now in a random temporal order without pairs.

## Methods

### Participants

We tested 8 patients (1 female; age range: 21-61; mean age = 37.8) who had been surgically implanted with intracranial electrodes for localization of seizure onset zone (see Table 1 for patient demographics and details on implant). This sample size was chosen a priori based on ***Sherman et al. (2022***). Two patients were tested a second time (two days later) because their first dataset was found to be unusable: one of these patients experienced severe eye irritation during the first testing session and there was a technical error with the triggers for the other patient. Electrode placement was determined solely by the clinical care team in order to localize seizure foci. Patients were recruited through the Yale Comprehensive Epilepsy Center and provided informed consent in a manner approved by the Yale University Human Subjects Committee. All data were collected at Yale New Haven Hospital.

**Table 1.** Patient demographics and electrode placement. Implant type indicates whether the implanted electrodes were only depth electrodes (Depth) or a combination of depth electrodes and grid/strip electrodes on the cortical surface (Combined). Hemisphere indicates the cerebral hemisphere into which the electrodes were implanted (see also Figure 2).

| Patient Information |  |  |  |  |  |
| --- | --- | --- | --- | --- | --- |
| ID | Age | Sex | nContacts | Implant Type | Hemisphere |
| 1 | 28 | M | 217 | Combined | Primarily Right |
| 2 | 21 | M | 119 | Combined | Left |
| 3 | 33 | M | 191 | Combined | Primarily Right |
| 4 | 34 | M | 116 | Combined | Right |
| 5 | 58 | M | 182 | Combined | Primarily Left |
| 6 | 44 | M | 168 | Combined | Left |
| 7 | 61 | F | 155 | Depth | Bilateral |
| 8 | 23 | M | 162 | Depth | Left |

### iEEG recordings

EEG data were recorded on a NATUS NeuroWorks EEG recording system. Data were collected at a sampling rate of 4096 Hz. Signals were referenced to an electrode chosen by the clinical team to minimize noise in the recording. To synchronize EEG signals with the experimental task, a custom-configured data acquisition system (DAQ) was used to convert signals from the research computer to 8-bit “triggers” that were inserted into a separate digital channel.

### iEEG preprocessing

iEEG preprocessing was carried out in FieldTrip (***Oostenveld et al., 2011***). A notch filter was applied to remove 60-Hz line noise. No re-referencing was applied. Data were downsampled to 256 Hz and segmented into trials using the triggers.

### Electrode localization

Electrode contact locations were identified using post-operative CT and MRI scans. Reconstructions were completed in BioImage Suite (***Papademetris et al., 2006***) and were subsequently registered to the patient’s pre-operative MRI scan, resulting in contact locations projected into the patient’s pre-operative space. The resulting files were converted from the Bioimagesuite format (.MGRID) into native space coordinates using FieldTrip functions. The coordinates were then used to create a mask in FSL (***Jenkinson et al., 2012***), with the coordinates of each contact occupying one voxel in the mask (Figure 2).

**Figure 2.**
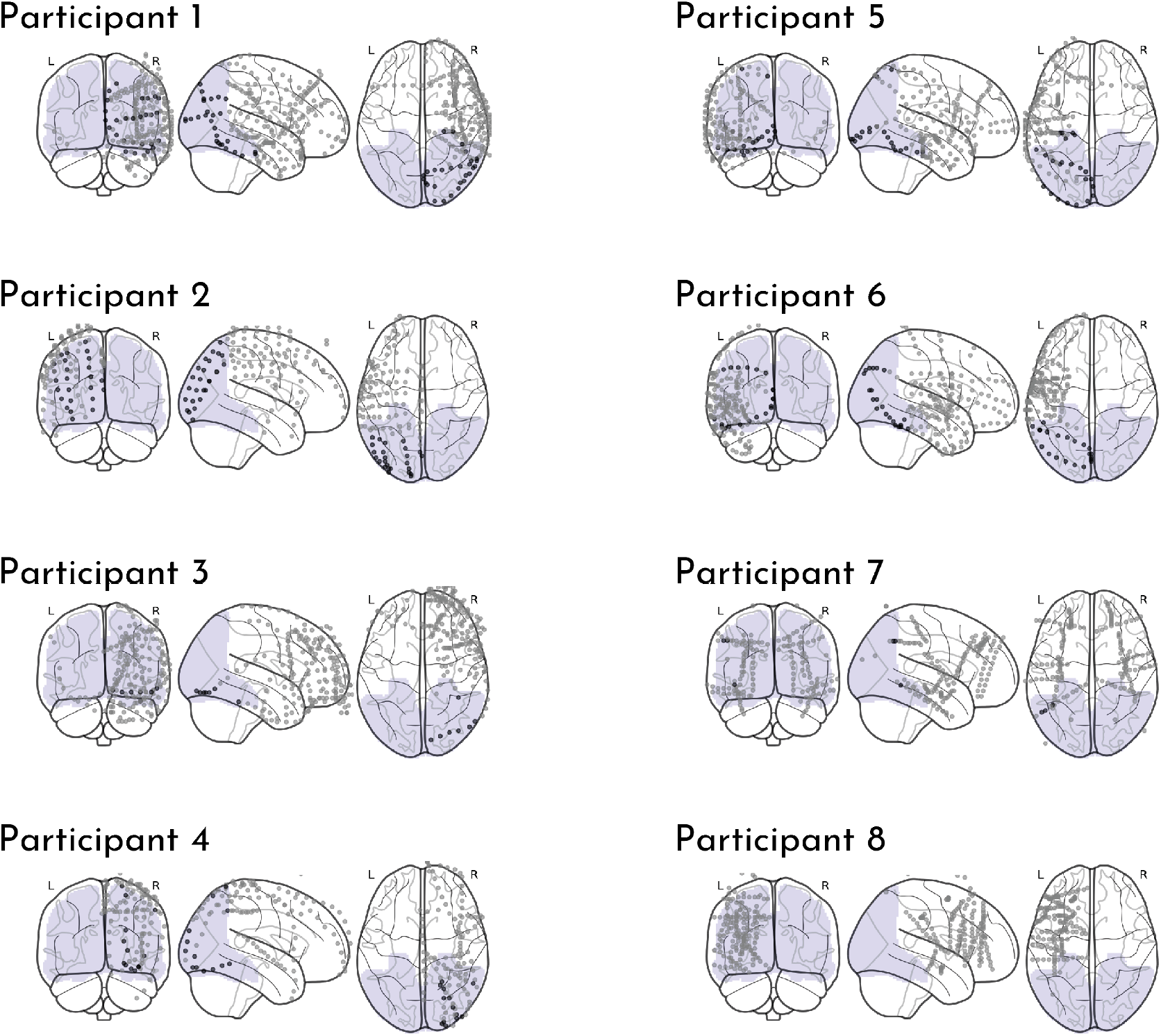
Electrode coverage for each patient. Each dot represents a single contact depicted on a standard glass brain. Contacts could be localized to the visual cortex ROI (purple shaded region) in 7 of the 8 patients, as indicated by darker black dots.

Given prior evidence for entrainment in sensory regions, we were interested in measuring neural responses in visual regions. We constructed a broad visual cortex region of interest (ROI), as in ***Sherman et al. (2022***), on the Montreal Neurological Institute (MNI) T1 2mm standard brain by combining the Occipital Lobe ROI from the MNI Structural Atlas and the following ROIs from the Harvard-Oxford Cortical Structural Atlas: Inferior Temporal Gyrus (temporo-ocipital part), Lateral Occipital Cortex (superior division), Lateral Occipital Cortex (inferior division), Intracalcarine Cortex, Cuneal Cortex, Parahippocampal Gyrus (posterior division), Lingual Gyrus, Temporal Occipital Fusiform Cortex, Occipital Fusiform Gyrus, Supracalcarine Cortex, and Occipital Pole. Each ROI was thresholded at 10% and then concatenated to create a single mask of visual cortex.

To localize contacts, we registered each patient’s pre-operative anatomical scan to the MNI T1 2mm standard brain template using linear registration (FSL FLIRT (***Jenkinson and Smith, 2001***; ***Jenkinson et al., 2002***)) with 12 degrees of freedom. We then used this registration matrix to transform each electrode mask into standard space. We overlaid the electrode masks onto the visual cortex ROI and onto the Harvard-Oxford cortical and subcortical structural atlases (maximum probability, 0 threshold). All but one of the patients had contacts in the visual cortex ROI, resulting in a final sample size of 7 participants for analyses of visual cortex.

### Stimuli

Task stimuli consisted of 720 unique scene images drawn from 18 distinct outdoor scene subcategories (amphitheater, amusement park, beach, bridge, canyon, desert, farm, forest, garden, highway, lake, lighthouse, marsh, mountain, park, sports field, town square, and waterfall; 40 images per subcategory). Six subcategories were randomly assigned to each of the Exemplar-level Structured, Category-level Structured, and Random conditions (see below). All images were collected from Google image searches and were cropped to a resolution of 600 × 800 pixels. Stimuli were presented using MATLAB with the Psychophysics toolbox (***Brainard, 1997***; ***Pelli, 1997***).

### Procedure

Participants completed the experiment on a laptop while seated in their hospital bed. The task consisted of at least one run of each of the three experimental conditions. During each run, participants passively viewed a rapid stream of scene images and were asked to pay attention to each image. To enable entrainment-based neural analyses, the stimulus-onset asynchrony (SOA) was fixed at 500 ms; each scene was presented for 250 ms, followed by a 250 ms inter-stimulus interval (ISI), during which a fixation cross appeared in the center of the screen. Each run sequence was 240 trials in length (2 mins of viewing time).

The Category-level Structured runs were our key runs of interest, in which we probed online abstraction of categorical regularities during statistical learning. Participants viewed a sequence of trial-unique scene images drawn from six scene categories. Participants were told in advance that they would be viewing images of scene categories and were given the names of the six categories. Unbeknownst to them, the six categories were assigned to three statistical pairs, such that a scene from one category (category A) was always followed by a scene from its paired category (category B; Figure 1, top right). Critically, these pairs existed only at the category-level because exemplars never repeated, requiring that patients abstract across exemplars in order to learn the regularities. No pair was allowed to repeat back-to-back in the sequence. In total, participants viewed 40 exemplars from each scene category once (40 repetitions of each category pair).

The Exemplar-level Structured runs served as a key comparison, enabling us to examine statistical learning of stimulus regularities without need for abstraction, as in prior studies (***Batterink and Paller, 2017***; ***Henin et al., 2021***). In this run, participants viewed a sequence containing multiple repetitions of six scene images, one each from six categories that did not overlap with the other conditions. Unbeknownst to them, the scenes were assigned to three statistical pairs (e.g., scene A -> scene B; Figure 1, middle right). No pair was allowed to repeat back-to-back in the sequence. Each exemplar/pair was repeated 40 times throughout the sequence.

The Random control runs served as our baseline condition, in which we did not expect any learning-related neural entrainment. As in the Exemplar-level Structured runs, participants viewed a sequence containing 40 repetitions of six scene images from six non-overlapping categories. In contrast to the two Structured conditions, the scenes were presented in a random order without reliable pairs that could be learned (Figure 1, bottom right). No individual scene was allowed to repeat within two images in the sequence.

Prior work has demonstrated that the order of statistical learning tasks can impact performance. Namely, learning is worse when one set of regularities is shown after another set or after randomness (***Jungé et al., 2007***; ***Gebhart et al., 2009***). Thus, to maximize our chance of detecting category-level neural entrainment, should it exist, especially given unexpected complications and interruptions in working with hospitalized patients, we tested the Category-level Structured condition first. We attempted to complete two of these runs back-to-back with the same sequence. When two runs were obtained (6/8 patients), we included data from both runs in all analyses. However, we also performed control analyses with only the first run, to equate the amount of data across conditions. After the Category-level Structured run(s), we completed one run of the Exemplar-level Structured and Random conditions next, counterbalancing order across participants. We decided on this semi-fixed condition order (Category-level Structured first) ahead of time, accepting that it could complicate comparison between conditions. However, note that each condition contains a positive control of neural entrainment to the individual image frequency, allowing us to assess data quality and ensure that conditions tested later in the session did not suffer from fatigue or inattention.

### Neural entrainment analyses

We conducted a phase coherence analysis to identify which electrode contacts entrained to our task. We examined entrainment at two frequencies: (1) the image frequency (2 Hz, corresponding to the 500-ms SOA between images), which reflects entrainment to the frequency of visual stimulation and should be present in all runs; and (2) the pair frequency (1 Hz, corresponding to the 1000-ms interval between pair onsets), which reflects entrainment to the statistical pairs and should only be present in the Structured runs (***Henin et al., 2021***).

For some runs of the task, there was a computer-based timing error such that the first trial’s ISI period was shorter than expected. Because the phase coherence analysis depends on reliable timing across trials, we excluded the first two trials from all analyses. The raw signals from the remaining 238 trials were segmented into 17 blocks comprising of 14 trials. For patients with two runs in the Category-level Structured condition, the raw signals were concatenated across runs, yielding 34 blocks.

We then converted the raw signals for each block into the frequency domain via fast Fourier transform and computed the phase coherence across blocks for each contact using the formula 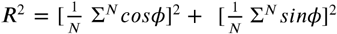, where *N* is the number of blocks and ϕ is the phase at a given frequency (***Ding and Simon, 2013; Henin et al., 2021***). Phase coherence was computed separately for each contact in the brain. We computed the peaks at the image and pair frequencies as the coherence at those frequencies relative to the coherence at the two neighboring frequencies (±0.14 Hz).

To assess statistical reliability across participants, we used a non-parametric, random-effects bootstrap resampling approach (***Efron and Tibshirani, 1986***). We first pooled the data across contacts and computed the effect of interest (e.g., mean or correlation coefficient). For each of 10,000 iterations, we randomly resampled the same sample size of participants with replacement (grabbing all of their electrodes) and recomputed the effect of interest to populate a sampling distribution of the effect. This sampling distribution was used to obtain 95% confidence intervals and perform null hypothesis testing. We calculated the *p*-value as the proportion of iterations in which the resampled effect had the opposite sign as the true effect; we then multiplied these values by 2 to obtain a two-tailed *p*-value. This tests the null hypothesis that the true effect is centered at zero and thus equally likely to be positive or negative by chance. A significant effect indicates that it did not matter which patients were resampled on any given iteration, and thus that the patients were interchangeable and the effect reliable across the sample. Across-participants resampling was performed in R (version 4.1.3), and the random number seed was set to 12345 before each resampling test.

To assess the reliability of a coherence peak *within* an individual electrode contact, we performed a randomization test. We shuffled the phase time series for each block 1,000 times, and recomputed the phase coherence across blocks of phase-shuffled data. We then computed the proportion of iterations that the true peak (coherence at the frequency of interest minus the neighboring frequencies) was larger than the null distribution of peaks to calculate the *p*-value. Given that we had a directional hypothesis (i.e., higher coherence than baseline), we did not multiply these *p* values by 2. Within-contact randomization testing was performed in MATLAB, and the random number seed was set to 12345 for each contact.

### Phase coherence timecourse analysis

To assess how neural entrainment to statistical pairs changed over the course of exposure, we performed a phase coherence timecourse analysis (***Henin et al., 2021***; ***Sherman et al., 2022***). We re-computed the coherence over an increasing number of blocks (e.g., first computing the coherence only between the first and second blocks, all the way up to all 17 blocks). For each cumulative block, we compared the coherence peak relative to a phase-shuffled surrogate dataset (described above) in order to compute the within-contact reliability. This resulted in a timecourse of p-values, allowing us to determine how many blocks of exposure were required for reliable entrainment. We performed this analysis at both the image and pair frequencies. We expected coherence at the image frequency to become reliable rapidly, as it reflects entrainment to sensory stimulation and does not require learning, providing a baseline for helping to interpret the timecourse of coherence at the (learned) pair frequency.

We computed p-value timecourses separately for the Category-level and Exemplar-level Structured conditions, focusing on visual contacts that showed reliable entrainment by the final block. That is, within each Structured condition, we averaged the timecourses of all contacts that exhibited reliable entrainment to the pair frequency by block 17. To equate opportunity for learning across patients, we only considered the first run of the Category-level Structured condition for patients with two of these runs.

We again assessed statistical reliability using a random-effects bootstrap resampling approach. We sought to quantify time to a significant response (number of cumulative blocks when *p* first went <0.05). To do so, we calculated the non-parametric *p*-value for a given number of cumulative blocks as the proportion of iterations in which the resampled p-value was less than 0.05. We then multiplied these values by 2 to obtain a two-tailed *p* value. This resampling test was done in R (version 4.1.3), with a random number seed of 12345.

## Results

### Evidence for category-level statistical learning in visual cortex

To assess whether the brain represents visual regularities online during learning, we capitalized on the fast, periodic nature of visual stimulation in our task and measured neural entrainment to the frequency of both individual images and statistical pairs (Figure 3A). Given prior work demonstrating neural entrainment in sensory regions (***Henin et al., 2021***; ***Sherman et al., 2022***), we focused our analyses on visual cortex. Specifically, we computed coherence within each contact localized to the visual cortex ROI (116 contacts) and averaged the coherence across contacts, within each participant. As a validation of our paradigm, we expected strong phase coherence at the frequency of image presentation in all three conditions. We further expected phase coherence at the frequency of pair presentation in the Exemplar-level Structured condition, replicating prior work demonstrating that the brain entrains to the frequency of statistical regularities (***Batterink and Paller, 2017***; ***Henin et al., 2021***). Critically, if the brain abstracts over these stimuli to learn higher-level, categorical regularities, we would expect phase coherence at the pair frequency in the Category-level Structured condition.

**Figure 3.**
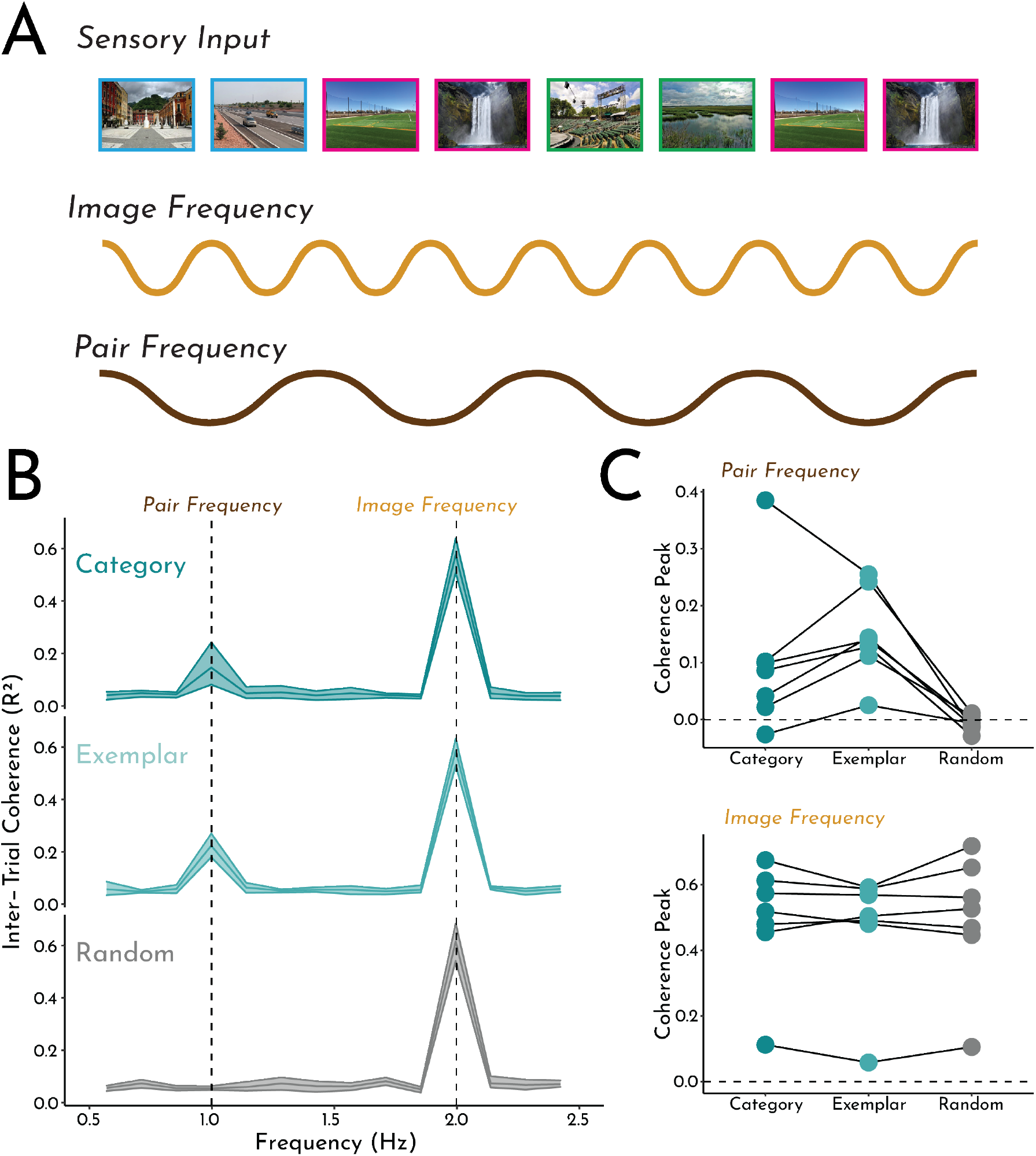
Phase coherence analysis. A) Schematic of analysis and hypothesized neural oscillations. We expected entrainment of visual contacts at the frequency of image presentation in all conditions. In the two Structured conditions (Exemplar-level and Category-level), we also expected entrainment at the frequency of (learned) pairs. B) These hypotheses were confirmed: We observed reliable peaks in coherence at the image frequency in all three conditions, but only at the pair frequency for the Category-level and Exemplar-level Structured conditions. Error shading indicates bootstrapped 95% confidence intervals. C) Coherence peaks at the pair frequency (top) and image frequency (bottom) for each participant across the three runs. Each circle/line represents one participant.

As shown in Figure 3B, we found reliable peaks in coherence at the image frequency in all three conditions (Exemplar-level Structured: mean difference, relative to neighboring frequencies = 0.53; 95% CI = [0.47, 0.57], *p* <0.001; Category-level Structured: mean difference = 0.54; 95% CI = [0.46, 0.60], *p* <0.001; Random: mean difference = 0.55; 95% CI = [0.47, 0.62], *p* <0.001). Critically, the peak in coherence at the pair frequency was reliable in both the Exemplar-level Structured condition (mean difference = 0.16, 95% CI = [0.12,0.21], *p* = 0.001) and the Category-level Structured condition (mean difference = 0.10, 95% CI = [0.041, 0.20], *p* <0.001), but not in the Random condition (mean difference = -0.0027, 95% CI = [-0.012, 0.0048], *p* = 0.50), providing online evidence for rapid statistical learning of exemplar pairs plus abstraction of category pairs.

To further understand these effects, we compared the peaks in coherence across conditions. We expected that there would be no condition differences in the peak at the image frequency, but that the peak at the pair frequency would be higher in Exemplar- and Category-level Structured conditions, relative to Random. Consistent with this hypothesis, there were no pairwise differences in the image frequency across conditions (Exemplar-level Structured vs. Random: mean difference = -0.026, 95% CI = [-0.066, 0.0094], *p* = 0.15; Category-level Structured vs. Random: mean difference = -0.018, 95% CI = [-0.047, 0.015], *p* = 0.28; Exemplar-vs. Category-level Structured: mean difference = -0.0073, 95% CI = [-0.042, 0.020], *p* = 0.61; Figure 3C, bottom). Importantly, the peak in coherence at the pair frequency was reliably higher for both Structured conditions than for the Random condition (Exemplar-level Structured vs. Random: mean difference = 0.17, 95% CI = [0.13, 0.21] *p* <0.001; Category-level Structured vs. Random: mean difference = 0.10, 95% CI = [0.043, 0.21], *p* <0.001; Figure 3C, top). Interestingly, the peak in coherence at the pair frequency was marginally higher in Exemplar-vs. Category-level Structured condition (mean difference = 0.062, 95% CI = [-0.0085, 0.11], *p* = 0.075), suggesting that stimulus regularities may be represented more robustly than abstract regularities in visual cortex, at least after a fixed and small amount of exposure.

The above analyses were performed on data concatenated across the two runs of the Category-level Structured condition (for participants with two runs). To confirm that evidence for categorical learning was not dependent on including more data, we repeated the analysis only considering the first Category-level Structured run. Indeed, we found a comparable peak in coherence at the pair frequency (mean difference = 0.096, 95% CI = [0.042, 0.19], *p* <0.001); the peak in coherence at the image frequency remained reliably high as well (mean difference = 0.56, 95% CI = [0.49, 0.62], *p* <0.001). Further, the peak in coherence at the pair frequency remained reliably higher than the Random condition (mean difference = 0.099, 95% CI = [0.044, 0.20], *p* <0.001) and marginally lower than that of the Exemplar-level condition (mean difference = -0.067, 95% CI = [-0.0028, 0.12], *p* = 0.061).

Together, these results demonstrate robust representation of statistical regularities in visual cortex, across levels of abstraction. After only two minutes of exposure, the visual cortex entrained not only to regularities at the exemplar-level (with the same exact image pairs repeating), but also to regularities that existed only at the category-level (requiring abstraction across exemplars in order to uncover the categorical structure). Critically, these data demonstrate that category-level regularities can be learned and represented *online* during learning, extending prior behavioral work which relied on delayed, offline test measures to infer that abstraction occurred.

### Co-representation of exemplar and category regularities

Above, we found evidence that visual cortex represents both exemplar- and category-level regularities. However, it is unclear whether these two effects are related. One possibility is that the more basic ability to extract regularities in sensory stimuli is a precursor for abstracting more complex regularities, in which case we might expect the same contacts to exhibit both effects and for the strength of these effects to be related. Another possibility is that stimulus learning and hierarchical abstraction are fundamentally distinct processes that may be implemented in different neural populations, and thus may be represented in different contacts and/or in the same contacts but in an unrelated manner.

To address this question, we first asked whether the strength of neural entrainment was correlated between conditions. Across all electrode contacts in the visual cortex ROI, we computed the Pearson correlation coefficient between the coherence peaks at the pair frequency. We found a reliable correlation in the pair frequency peak for Category- and Exemplar-level Structured conditions (*r* = 0.33, 95% CI = [0.019, 0.58], *p* = 0.033; Figure 4A). In contrast, there was no reliable correlation between the Random condition and either the Category-level Structured condition (*r* = -0.10, 95% CI = [-0.24, 0.077], *p* = 0.22) or the Exemplar-level Structured condition (*r* = -0.011, 95% CI = [-0.14, 0.14], *p* = 0.82). The modest correlation between coherence for exemplar pairs and category pairs suggests a degree of shared representation of regularities across levels of abstraction. Importantly, given that we did not find such correlations with the Random condition, we can be confident that this correlation was not driven by generic across-contact factors such as baseline coherence or data quality.

**Figure 4.**
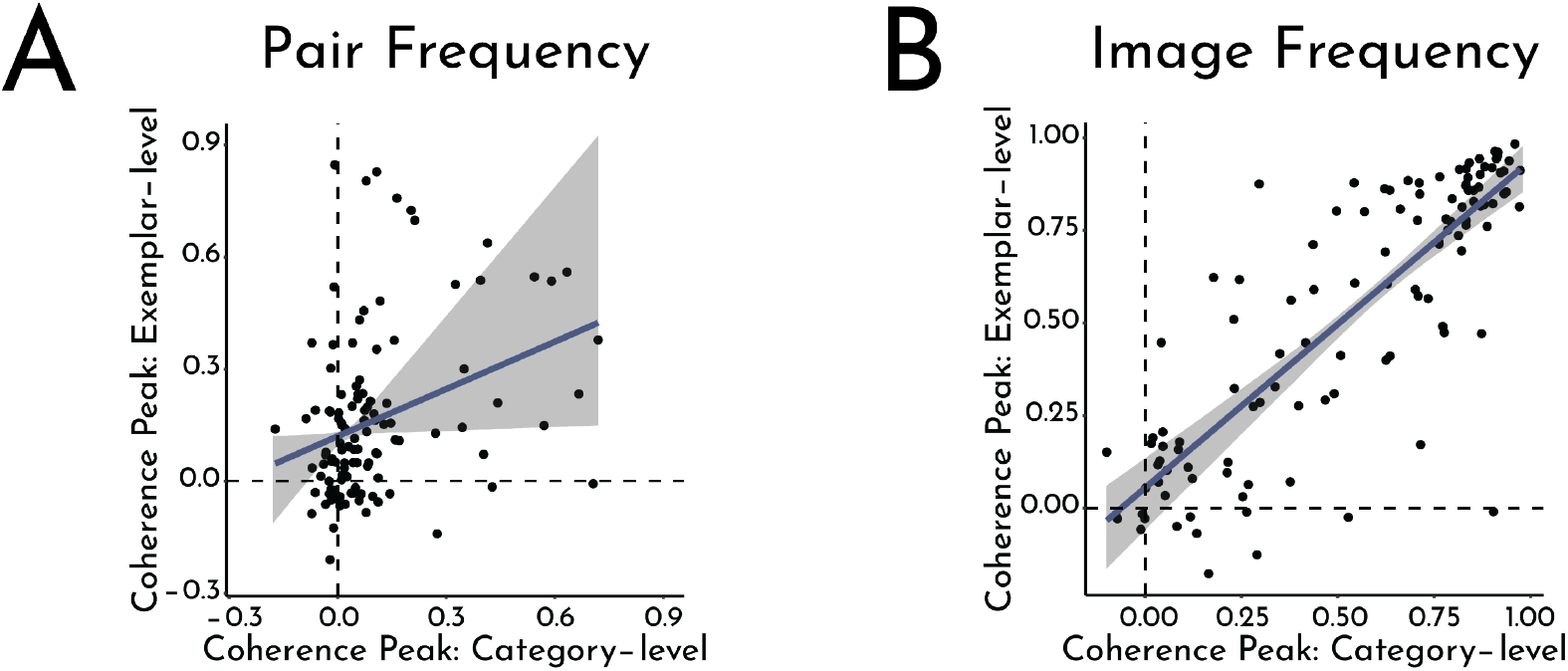
Correlations across contacts. A) Correlation between the coherence peak at the pair frequency in the Category-level Structured condition and the coherence peak at the pair frequency in the Exemplar-level Structured condition. B) Correlation between the coherence peak at the image frequency in the Category-level Structured condition and the coherence peak at the image frequency in the Exemplar-level Structured condition. Each circle represents an electrode contact. Error shading indicates bootstrapped 95% confidence intervals.

As a further control, we computed the pairwise correlations for the image frequency peaks. Unlike the pair frequency, we did not expect these correlations to differ between conditions. Indeed, we found high correlations across the board (Category- and Exemplar-level Structured, Figure 4B: *r* = 0.83, 95% CI = [0.70, 0.92], *p* <0.001; Category-level Structured and Random: *r* = 0.88, 95% CI = [0.81, 0.93], *p* <0.001; Exemplar-level Structured and Random: *r* = 0.87, 95% CI = [0.79, 0.93], *p* <0.001).

To further address the relationship between exemplar and category regularities, we labeled individual contacts according to whether they exhibited a reliable coherence peak at the frequencies of interest in each condition. Of the 116 total electrode contacts in visual cortex, 67 exhibited entrainment to the pair frequency in one or both Structured conditions; 27 entrained to the pair frequency in the Exemplar-level Structured condition only, 12 in the Category-level Structured condition only, and 28 in both Structured conditions. To assess whether this is more overlap than would be expected by chance, given the number of reliable contacts in each condition, we independently shuffled the correspondence between contacts and significance labels across conditions and recomputed the overlap. We found that the observed overlap was indeed reliable (mean null overlap = 19 contacts, 95% CI = [14, 24], *p* <0.001), indicating that some parts of visual cortex exhibit a dual representation both exemplar and category regularities.

To understand whether these dual-coding contacts were responsible for the correlations observed above, we re-computed the correlations after removing these contacts. Indeed, this eliminated the correlation (88 non-overlapping contacts: *r* = -0.086, 95% CI = [-0.18, 0.15], *p* = 0.31). However, there was also no reliable correlation when restricting the analysis to only the dual-coding contacts (28 overlapping contacts: *r* = 0.041, 95% CI = [-0.36, 0.33], *p* = 0.41). This suggests that the original correlation benefitted from variance in coding properties across contacts and/or from the greater sensitivity provided by a larger sample size of contacts.

### Examining the timecourse of learning in visual cortex

We have presented evidence that populations of electrode contacts in visual cortex entrain to both exemplar- and category-level regularities online during statistical learning. However, it is possible that statistical learning of more abstract category regularities requires more exposure than learning of simpler, stimulus-driven exemplar regularities. To assess the evolution of entrainment over the course of learning and whether it differs across conditions, we performed a timecourse analysis. Specifically, we re-computed coherence over an increasing number of blocks (e.g., first computing the coherence only between the first and second blocks, then between the first, second and third blocks, all the way up to 17 blocks) to determine the block count at which contacts exhibited reliable entrainment. In other words, we asked how much exposure was required for contacts that exhibited reliable entrainment in the final block to reach a statistically reliable response. For the Category-level Structured condition, we only analyzed each patient’s first run in order to equate the opportunity for learning both across patients and conditions.

In the Category-level Structured condition (Figure 5A, left), we found reliable entrainment only when computing coherence across 16 or more blocks (16 blocks: mean *p* = 0.023, 95% CI = [0.0033, 0.044], *p* = 0.0056; 17 blocks: mean *p* = 0.0097, 95% CI = [0.0023, 0.018], *p* <0.001). In the Exemplar-level Structured condition (Figure 5A, right), entrainment appeared marginally after 14 blocks (mean *p* = 0.029, 95% CI = [0.011, 0.052], *p* = 0.057) and reliably for 15 or more blocks (*p*s <0.001). These data suggest that exemplar and category regularities were learned at a similar timescale, with slightly faster acquisition for exemplar regularities.

**Figure 5.**
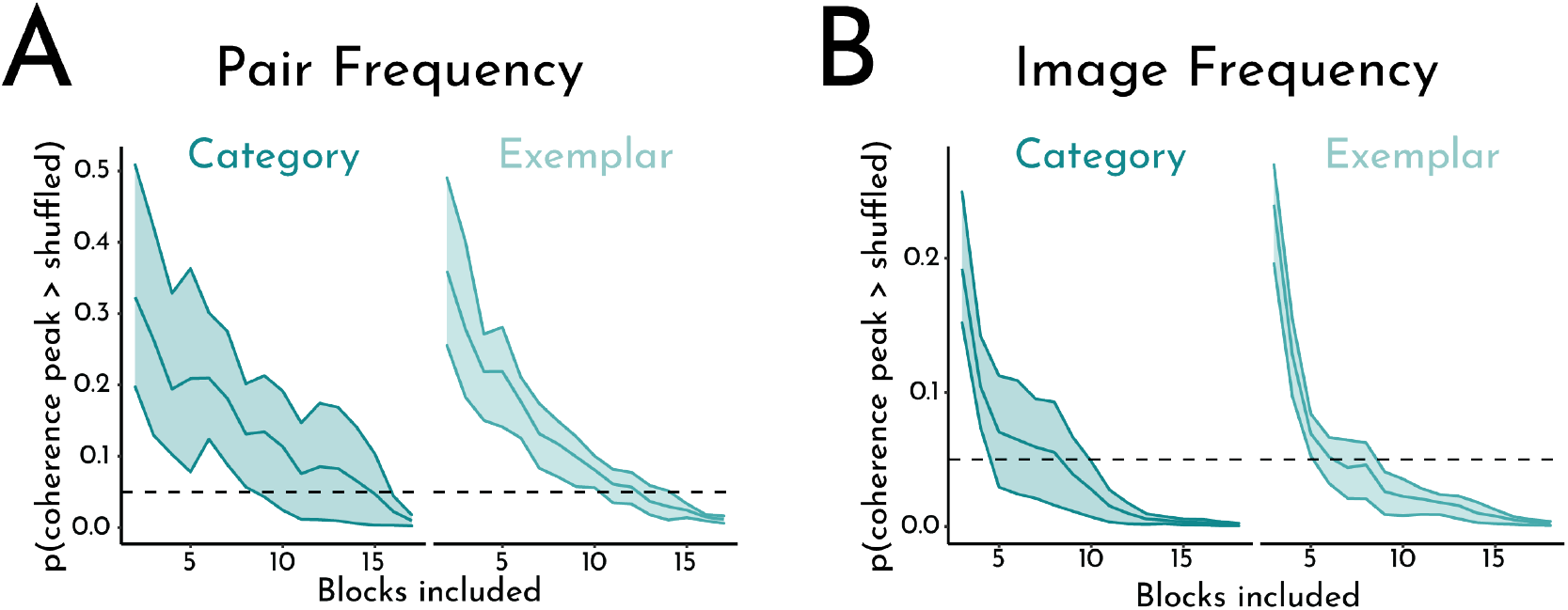
Timecourse analysis. A) Emergence of a reliable phase coherence peak at the pair frequency across blocks in the Category-level Structured (left) and Exemplar-level Structured (right) conditions. For each cumulative block count N, we computed the proportion of iterations that the coherence peak across N blocks was greater than the peak across N blocks of phase-shuffled data to obtain a p-value; we then determined the first block at which the permuted p-value across contacts was reliably less than 0.05 (dashed line). B) Emergence of a significant response at the image frequency across blocks in the Category-level Structured (left) and Exemplar-level Structured (right) conditions. Error shading indicates bootstrapped 95% confidence intervals.

To establish a floor of how quickly we might theoretically expect to see a reliable entrainment effect, we performed this same analysis for the image frequency (again, only considering contacts that exhibited reliable entrainment to the image frequency in the final block). Because entrainment to the images was given by the sensory input and not from statistical learning, we did not expect meaningful differences between conditions. In the Category-level Structured condition (Figure 5B, left), there was reliable coherence at the image frequency by block 9 (mean *p* = 0.028, 95% CI = [0.0072, 0.049], *p* = 0.037; all subsequent block counts, *p*s <0.001). The Exemplar-level Structured condition (Figure 5B, right) followed a similar pattern, with reliable entrainment by block 8 (mean *p* = 0.026, 95% CI = [0.009, 0.041], *p* = 0.0018; all subsequent block counts, *p*s <0.001). Finally, we also computed the timecourse of the image frequency effect in the Random condition and found a similar pattern, with reliable entrainment by block 9 (mean *p* = 0.033; 95% CI = [0.016, 0.048], *p* = 0.024; block 10: mean *p* = 0.034, 95% CI = [0.016, 0.049], *p* = 0.036; all subsequent block counts, *p*s <0.001).

### Categorical abstraction during statistical learning across the brain

We initially focused on how visual cortex represents visual regularities given our prior work (***Sherman et al., 2022***), but a wide range of brain regions have been implicated in statistical learning (***Batterink et al., 2019***; ***Henin et al., 2021***). To examine online abstraction during statistical learning more broadly, we measured neural entrainment to exemplar and category regularities in an exploratory brain-wide analysis.

First, as in the analysis restricted to visual cortex, we identified which contacts represented exemplar and/or category regularities by testing for reliable phase coherence at the pair frequency relative to neighboring frequencies. Of a total of 1,310 contacts across all patients, we found reliable entrainment at the pair frequency in 175 contacts for the Exemplar-level Structured condition and in 177 contacts for the Category-level Structured condition; 41 of these contacts overlapped. This amount of overlap was reliably greater than expected by chance (Figure 6A; mean null overlap = 24 contacts, 95% CI = [16, 32], *p* <0.001). Because this brain-wide analysis included visual cortex, it is possible that the reliable overlap was driven by visual contacts, which we earlier showed exhibited reliable overlap. We therefore repeated the brain-wide analysis after excluding contacts in the visual cortex ROI. Of the remaining 1,194 contacts across all patients, we found reliable entrainment at the pair frequency in 120 contacts for the Exemplar-level Structured condition and 137 contacts for the Category-level Structured condition; 13 of these contacts overlapped. However, this amount of overlap was not reliably greater than what would be expected by chance (mean null overlap = 14 contacts, 95% CI = [8, 20], *p* = 0.52), suggesting that dual coding of exemplar- and category-level regularities in individual contacts was restricted to visual cortex.

**Figure 6.**
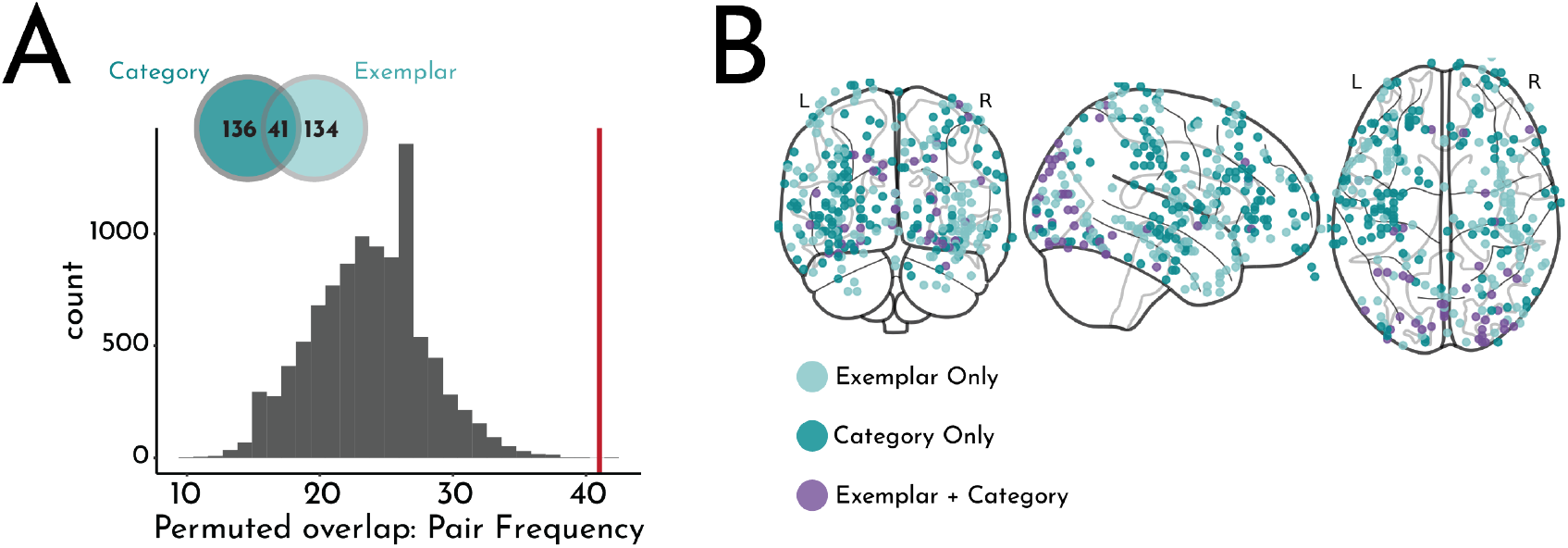
Exploratory brain-wide analyses. A) Histogram: distribution of how many contacts would be expected to entrain to the pair frequency in *both* the Exemplar- and Category-level Structured conditions by chance; red line indicates the observed overlap, indication that many more contacts coded for both exemplar and category regularities than would be expected by chance. Inset: Venn diagram illustrating the total number of contacts that entrained to the pair frequency in both conditions and their overlap. B) Map of contacts (across all patients) that entrained to the pair frequency in one or both conditions on a standard glass brain.

We next sought to localize these structure-sensitive contacts throughout the brain (Figure 6B). We mapped the contacts onto the Harvard-Oxford cortical and subcortical atlases and quantified how many contacts exhibited effects within each gray-matter atlas ROI. Table 2 summarizes the results by listing atlas ROIs that contained at least 5 contacts that entrained at an uncorrected level to the pair frequency in at least one of the Structured conditions. Consistent with our planned visual ROI, many of these contacts were located in visual cortex (e.g., lateral occipital cortex, lingual gyrus, occipital pole). However, we also observed entrainment to learned regularities in frontal and anterior temporal regions, some showing a preference for regularities available directly in the exemplar stimuli (e.g., temporal pole) and others for regularities that required categorical abstraction (e.g., frontal pole and precentral gyrus). Importantly, claims about localization in the brain are limited by the fact that we did not have full coverage of all brain regions, given that electrode placement was determined clinically.

**Table 2.**
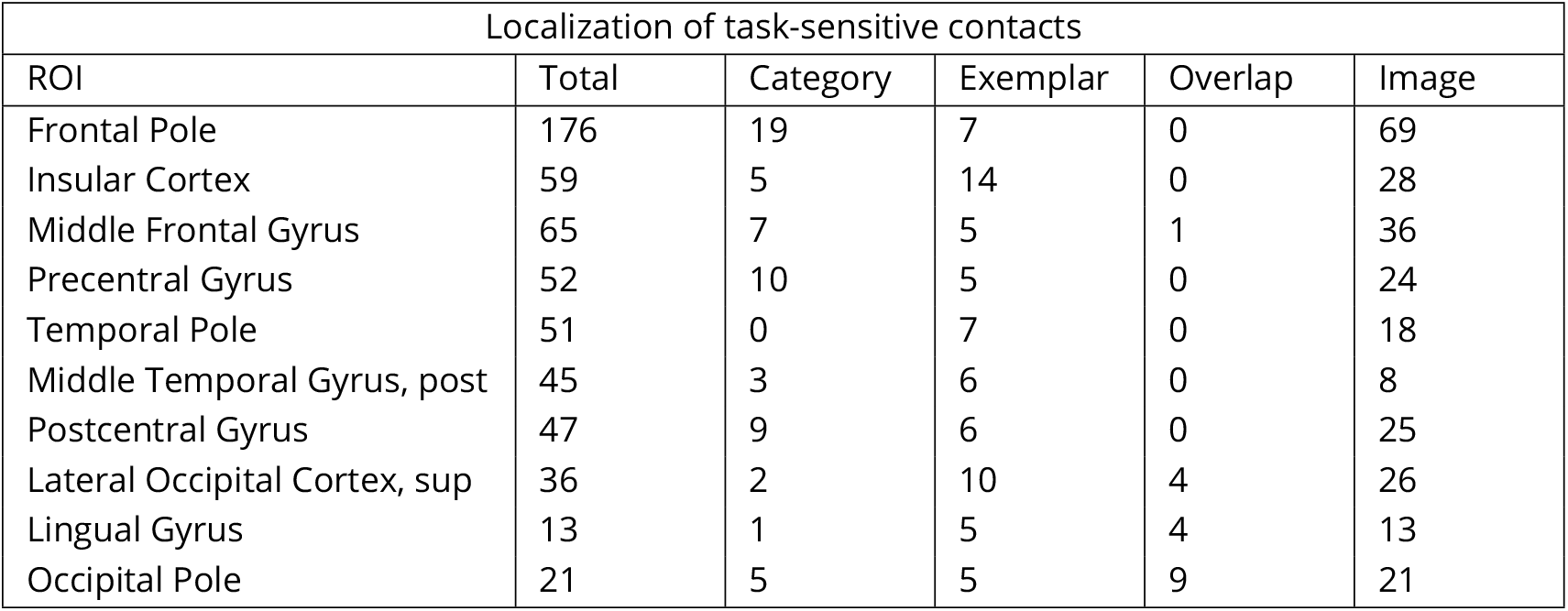
Gray-matter ROIs in the Harvard-Oxford cortical and subcortical atlases that contained at least 5 contacts with reliable entrainment at the pair frequency in the Category- and/or Exemplar-level Structured conditions. We also included the total number of contacts in each ROI (Total) and the number of contacts that entrained at the image frequency in the Category- and/or Exemplar-level Structured conditions (Image).

## Discussion

In the current study, we capitalized on the high spatial and temporal precision of intracranial EEG to explore how the brain learns and represents statistical regularities across varying levels of abstraction. Specifically, we contrasted the learning of exemplar-level regularities (defined by the transition probabilities between individual images) with the learning of category-level regularities (defined by the transition probabilities between image categories, thus requiring abstraction across individual images). We found robust representation of both kinds of regularities in visual cortex and throughout the brain during statistical learning. These findings speak to several issues in the statistical learning literature and raise questions for future research.

### Online evidence for category-level statistical learning

In measuring neural entrainment to the frequency of regularities, we employed a covert, online measure of statistical learning. This builds on a body of work that measured category-level statistical learning with offline behavioral tests, such as asking participants to judge their familiarity with pairs of images or categories (***Brady and Oliva, 2008***; ***Otsuka et al., 2013***; ***Emberson and Rubinstein, 2016***; ***Jung et al., 2021***). However, it is unclear whether above-chance performance on these tests reflects abstraction of category relationships during the learning process itself or the formation of specific stimulus associations during learning that enabled successful inferences about test items from the same categories. It is also possible that these behavioral studies engendered both abstraction during learning and inferences at test, yet it remains unclear which effect (or both) drove test performance. Further complicating the interpretation of offline behavioral performance as evidence of online abstraction, online and offline measures of statistical learning are not always correlated (***Kiai and Melloni, 2021***). The current study sought to skirt these interpretational challenges by measuring neural entrainment as an online neural index of statistical learning. The observed entrainment to category pairs provides novel evidence for rapid statistical learning between abstractions over individual exemplars.

One limitation of our study is that it is unclear how the neural entrainment measure of statistical learning and abstraction relates to more canonical behavioral measures. Given our short testing time with each patient, their limited energy and attention span, and the small number of patients, we optimized our task design and testing time for neural rather than behavioral measures of learning. Future studies could perhaps use scalp EEG in a well-powered normative sample to help link neural and behavioral measures of category-level statistical learning. Future studies could further consider how neural entrainment during learning relates to both online (e.g., response time) and offline (e.g., familiarity) behavioral measures; that said, it may be difficult to develop online behavioral measures during a task designed for neural entrainment, given the fast presentation rates that such tasks require. Prior studies have demonstrated that neural evidence of statistical learning can appear earlier and even in the absence of behavioral evidence of learning (***Turk-Browne et al., 2009***); thus, it is possible that our current results reflect a rapid sensitivity of the brain to category regularities.

Additional limitations apply in how to interpret the timecourse results. Although these results provide evidence that learning occurs quite quickly (less than two minutes) in both Structured conditions, it is unclear how this maps onto the underlying trajectory of learning. We found reliable evidence of statistical learning for exemplar regularities two blocks earlier than for category regularities. Does this small difference in the amount of required exposure mean that specific stimulus associations must be learned before more abstract associations? Or perhaps the individual images were represented both as exemplars and categories during perception and associations were learned at both levels in parallel? In this case, learning of category regularities may be slower because of the added complexity in dealing with greater input variability (e.g., in the extent to which a given exemplar was a prototype of a category). Note also that the “time to significant response” measure we used based on prior work (***Henin et al., 2021***) is relatively conservative and constrained (measuring the reliability of responses within each contact) and does not necessarily reflect the veridical overall timecourse of learning across the brain or in behavior. Further, we computed this metric by averaging across contacts that were reliable in the final block, which may have obscured heterogeneous timecourses for different aspects or stages of learning across the brain.

Finally, different aspects of learned structure can be measured. For example, memory for the temporal order of items within a statistical unit (e.g., triplet) can be dissociated from memory for the item groupings (***Park et al., 2018***; ***Forest et al., 2022***), and these distinct types of memory may be supported by different underlying neural representations (***Davachi and DuBrow, 2015***; ***Henin et al., 2021***). Although providing evidence of learning overall, the current study, and the basic neural entrainment design it employed, is insensitive to these differing underlying representations. Future studies could employ other neural measures, such as pre- and post-learning templates (***Schapiro et al., 2012***), to assess changes in the representations of the individual paired items. Such measures could be used to test hypotheses about how these constituent items are represented at different levels of abstraction as a function of statistical learning.

### Local and distributed representations of visual regularities across the brain

We focused our main analyses on visual cortex, which we hypothesized would show neural entrainment to visual regularities between visual images (***Henin et al., 2021***; ***Sherman et al., 2022***). However, we also performed an exploratory brain-wide analysis to uncover where category and exemplar regularities were represented throughout the brain. This analysis largely confirmed our a priori choice to focus on visual cortex, but also revealed a distributed representation of structure, with many frontal (e.g., frontal pole, insula, middle frontal gyrus, and precentral gyrus) and temporal (e.g., temporal pole, middle temporal gyrus) regions also exhibiting entrainment to visual regularities. These findings are largely consistent with prior fMRI studies demonstrating sensitivity to structure in these regions (***Turk-Browne et al., 2009, 2010***; ***Karuza et al., 2013, 2017***).

This analysis revealed relatively little evidence that entire brain regions specialize at a particular level of abstraction. Although some regions exhibited a bias towards one level (e.g., more contacts in the frontal pole entrained only to category regularities, and more contacts in the insula entrained only to exemplar regularities), very few regions solely represented one level. The only exception was the temporal pole, which only exhibited entrainment to exemplar-level regularities. Similarly, most contacts did not show a general sensitivity to structure regardless of abstraction. The small (but reliable) number of such contacts representing *both* category and exemplar regularities were restricted to visual cortex (e.g., occipital pole). Still, the majority of visual contacts entrained to one level of structure or the other, but not both. At the level of entire brain regions, some regions contained distinct contacts that entrained selectively to category and exemplar regularities, yet no contacts that entrained to both. This raises the possibility that there may be distinct neural populations and cognitive processes even within the same brain region for statistical learning at varying levels of abstraction.

An important limitation to these exploratory brain-wide analyses is that they only had access to partial coverage of the brain. Although we had relatively broad coverage of cortical regions for an intracranial EEG study, the electrode locations were chosen entirely for clinical purposes and were thus not always comprehensive or standardized across patients. However, this is an expected limitation for any iEEG based study. Further, we had insufficient coverage of the hippocampus in this sample (only 5 contacts across all patients), a region that has been consistently implicated in rapid statistical learning (***Turk-Browne et al., 2009***; ***Schapiro et al., 2012***; ***Covington et al., 2018***; ***Sherman and Turk-Browne, 2020***; ***Henin et al., 2021***; ***Graves et al., 2022***). Future studies could recruit a more targeted sample of intracranial EEG patients (e.g., with hippocampal depth electrodes) or use fMRI for high-resolution hippocampal coverage potentially across a larger sample of individuals.

## Conclusions

Together, our results provide evidence for rapid and robust online abstraction of categorical regularities during statistical learning. This occurred heavily within visual cortex, suggesting a remarkable capability for the brain to aggregate across noisy, idiosyncratic instances to extract stable properties of the environment that can generalize to new situations.

## Acknowledgments

We are grateful to the patients who participated in the study and to the physicians, fellows, and support staff who cared for them at the Yale New Haven Hospital. We thank Kun Wu for providing the electrode reconstructions; Simon Henin for advice on statistical analyses; and Erica Busch for advice on the brain-wide analyses. This project was supported by a National Science Foundation Graduate Research Fellowship (B.E.S.) and the Canadian Institute for Advanced Research (N.B.T-B.).

## References

Batterink, L. (2020). Syllables in sync form a link: Neural phase-locking reflects word knowledge during language learning. Journal of Cognitive Neuroscience, 32(9):1735–1748.

Batterink, L. J. and Paller, K. A. (2017). Online neural monitoring of statistical learning. Cortex, 90:31–45.

Batterink, L. J., Paller, K. A., and Reber, P. J. (2019). Understanding the neural bases of implicit and statistical learning. Topics in Cognitive Science, 11(3):482–503.

Bauer, A.-K. R., Debener, S., and Nobre, A. C. (2020). Synchronisation of neural oscillations and cross-modal influences. Trends in Cognitive Sciences, 24(6):481–495.

Brady, T. F. and Oliva, A. (2008). Statistical learning using real-world scenes: Extracting categorical regularities without conscious intent. Psychological Science, 19(7):678–685.

Brainard, D. H. (1997). The psychophysics toolbox. Spatial Vision, 10(4):433–436.

Choi, D., Batterink, L. J., Black, A. K., Paller, K. A., and Werker, J. F. (2020). Preverbal infants discover statistical word patterns at similar rates as adults: Evidence from neural entrainment. Psychological Science, 31(9):1161–1173.

Covington, N. V., Brown-Schmidt, S., and Duff, M. C. (2018). The necessity of the hippocampus for statistical learning. Journal of Cognitive Neuroscience, 30(5):680–697.

Davachi, L. and DuBrow, S. (2015). How the hippocampus preserves order: the role of prediction and context. Trends in Cognitive Sciences, 19(2):92–99.

Ding, N., Melloni, L., Zhang, H., Tian, X., and Poeppel, D. (2016). Cortical tracking of hierarchical linguistic structures in connected speech. Nature Neuroscience, 19(1):158–164.

Ding, N. and Simon, J. Z. (2013). Power and phase properties of oscillatory neural responses in the presence of background activity. Journal of computational neuroscience, 34(2):337–343.

Efron, B. and Tibshirani, R. (1986). Bootstrap methods for standard errors, confidence intervals, and other measures of statistical accuracy. Statistical Science, pages 54–75.

Emberson, L. L. and Rubinstein, D. Y. (2016). Statistical learning is constrained to less abstract patterns in complex sensory input (but not the least). Cognition, 153:63–78.

Forest, T. A., Finn, A. S., and Schlichting, M. L. (2022). General precedes specific in memory representations for structured experience. Journal of Experimental Psychology: General, 151(4):837.

Gebhart, A. L., Aslin, R. N., and Newport, E. L. (2009). Changing structures in midstream: Learning along the statistical garden path. Cognitive Science, 33(6):1087–1116.

Graves, K. N., Sherman, B. E., Huberdeau, D., Damisah, E., Quraishi, I. H., and Turk-Browne, N. B. (2022). Remembering the pattern: A longitudinal case study on statistical learning in spatial navigation and memory consolidation. Neuropsychologia, 174:108341.

Henin, S., Turk-Browne, N. B., Friedman, D., Liu, A., Dugan, P., Flinker, A., Doyle, W., Devinsky, O., and Melloni, L. (2021). Learning hierarchical sequence representations across human cortex and hippocampus. Science Advances, 7(8):eabc4530.

Jenkinson, M., Bannister, P., Brady, M., and Smith, S. (2002). Improved optimization for the robust and accurate linear registration and motion correction of brain images. NeuroImage, 17(2):825–841.

Jenkinson, M., Beckmann, C. F., Behrens, T. E., Woolrich, M. W., and Smith, S. M. (2012). Fsl. NeuroImage, 62(2):782–790.

Jenkinson, M. and Smith, S. (2001). A global optimisation method for robust affine registration of brain images. Medical Image Analysis, 5(2):143–156.

Jun, J. and Chong, S. C. (2018). Visual statistical learning at basic and subordinate category levels in real-world images. Attention, Perception, & Psychophysics, 80(8):1946–1961.

Jung, Y., Walther, D. B., and Finn, A. S. (2021). Children automatically abstract categorical regularities during statistical learning. Developmental Science, 24(5):e13072.

Jungé, J. A., Scholl, B. J., and Chun, M. M. (2007). How is spatial context learning integrated over signal versus noise? a primacy effect in contextual cueing. Visual Cognition, 15(1):1–11.

Karuza, E. A., Emberson, L. L., Roser, M. E., Cole, D., Aslin, R. N., and Fiser, J. (2017). Neural signatures of spatial statistical learning: Characterizing the extraction of structure from complex visual scenes. Journal of Cognitive Neuroscience, 29(12):1963–1976.

Karuza, E. A., Newport, E. L., Aslin, R. N., Starling, S. J., Tivarus, M. E., and Bavelier, D. (2013). The neural correlates of statistical learning in a word segmentation task: An fmri study. Brain and Language, 127(1):46–54.

Kiai, A. and Melloni, L. (2021). What canonical online and offline measures of statistical learning can and cannot tell us. bioRxiv.

Luo, Y. and Zhao, J. (2018). Statistical learning creates novel object associations via transitive relations. Psychological science, 29(8):1207–1220.

Norcia, A. M., Appelbaum, L. G., Ales, J. M., Cottereau, B. R., and Rossion, B. (2015). The steady-state visual evoked potential in vision research: A review. Journal of Vision, 15(6):4–4.

Nozaradan, S., Peretz, I., Missal, M., and Mouraux, A. (2011). Tagging the neuronal entrainment to beat and meter. Journal of Neuroscience, 31(28):10234–10240.

Oostenveld, R., Fries, P., Maris, E., and Schoffelen, J.-M. (2011). Fieldtrip: open source software for advanced analysis of meg, eeg, and invasive electrophysiological data. Computational Intelligence and Neuroscience, 2011.

Otsuka, S., Nishiyama, M., Nakahara, F., and Kawaguchi, J. (2013). Visual statistical learning based on the perceptual and semantic information of objects. Journal of Experimental Psychology: Learning, Memory, and Cognition, 39(1):196.

Papademetris, X., Jackowski, M. P., Rajeevan, N., DiStasio, M., Okuda, H., Constable, R. T., and Staib, L. H. (2006). Bioimage suite: An integrated medical image analysis suite: An update. The Insight Journal, 2006:209.

Park, S. H., Rogers, L. L., and Vickery, T. J. (2018). The roles of order, distance, and interstitial items in temporal visual statistical learning. Attention, Perception, & Psychophysics, 80(6):1409–1419.

Pelli, D. G. (1997). The videotoolbox software for visual psychophysics: transforming numbers into movies. Spatial Vision.

Preston, A. R. and Eichenbaum, H. (2013). Interplay of hippocampus and prefrontal cortex in memory. Current Biology, 23(17):R764–R773.

Schapiro, A. C., Kustner, L. V., and Turk-Browne, N. B. (2012). Shaping of object representations in the human medial temporal lobe based on temporal regularities. Current Biology, 22(17):1622–1627.

Sherman, B. E., Graves, K. N., Huberdeau, D. M., Quraishi, I. H., Damisah, E. C., and Turk-Browne, N. B. (2022). Temporal dynamics of competition between statistical learning and episodic memory in intracranial recordings of human visual cortex. Journal of Neuroscience, 42(48):9053–9068.

Sherman, B. E., Graves, K. N., and Turk-Browne, N. B. (2020). The prevalence and importance of statistical learning in human cognition and behavior. Current Opinion in Behavioral Sciences, 32:15–20.

Sherman, B. E. and Turk-Browne, N. B. (2020). Statistical prediction of the future impairs episodic encoding of the present. Proceedings of the National Academy of Sciences, 117(37):22760–22770.

Störmer, V. S. and Alvarez, G. A. (2014). Feature-based attention elicits surround suppression in feature space. Current Biology, 24(17):1985–1988.

Turk-Browne, N. B., Scholl, B. J., Chun, M. M., and Johnson, M. K. (2009). Neural evidence of statistical learning: Efficient detection of visual regularities without awareness. Journal of Cognitive Neuroscience, 21(10):1934–1945.

Turk-Browne, N. B., Scholl, B. J., Johnson, M. K., and Chun, M. M. (2010). Implicit perceptual anticipation triggered by statistical learning. Journal of Neuroscience, 30(33):11177–11187.

Zhou, Z., Singh, D., Tandoc, M. C., and Schapiro, A. C. (2021). Distributed representations for human inference. bioRxiv.

